# *cctbx.xfel*: a suite for processing serial crystallographic data

**DOI:** 10.1101/2025.05.04.652045

**Authors:** Aaron S. Brewster, Daniel W. Paley, Asmit Bhowmick, David W. Mittan-Moreau, Iris D. Young, Derek A. Mendez, Daniel M. Tchoń, Billy K. Poon, Nicholas K. Sauter

## Abstract

The *cctbx.xfel* suite of processing programs and tools allows fast, visual analysis of serial diffraction images from synchrotrons and XFELs. Built on *DIALS* and *cctbx*, *cctbx.xfel* is designed for real-time and post-experiment processing with a fully featured graphical user interface. Users can quickly identify hitrates, view diffraction patterns, analyze unit-cell isomorphism using clustering, and merge data using a metadata tagging approach that allows on-the-fly organization and visualization of processing results. This paper describes the fundamental algorithms and command-line programs used by *cctbx.xfel*, including the two main program *dials.stills process*, which performs spot-finding, indexing, geometric refinement, and integration, and *cctbx.xfel.merge*, which performs scaling, post-refinement, and merging. A discussion of merging statistics is presented and newer features are described, including random sub-sampling for indexing multi-lattice hits and Δ*CC*_1_*_/_*_2_ filtering to remove outliers. Finally we show a complex, heterogeneous sample containing hexagonal and monoclinic isoforms in *P* 6_3_ and *P* 2_1_. The isoforms are separated by unit cell clustering, and for each isoform we resolve a (pseudo-)merohedral indexing ambiguity.

## 1. Introduction

Serial crystallography experiments at X-ray free electron lasers (XFELs) and synchrotrons can produce 10^2^ to 10^6^ protein diffraction images at collection rates from 10 Hz to 3-5 kHz. Diffraction patterns need to be identified and analyzed as fast as possible to enable fast feedback to users and operators, and specialized algorithms need to be used to handle still images, where every pattern arises from a distinct crystal in a random orientation. For fast and accurate processing of serial diffraction data, we developed *cctbx.xfel*, a suite of serial crystallographic data reduction programs built on the packages *cctbx* (Grosse-Kunstleve *et al*., 2002; Sauter *et al*., 2013) and *DIALS* (Winter *et al*., 2018). In this paper, we will describe the algorithms and most common options used for the two main programs, *dials.stills process*, which models data on individual diffraction patterns (§2), and *cctbx.xfel.merge*, which is responsible for the global merging of duplicate measurements on all the diffraction patterns (§4). We treat issues that are unique to serial crystallography, such as the slow drift of instrumentation (§3) and the recruitment of sufficient computational infrastructure to enable fast processing (§5). We show how a new option, random sub-sampling, enables more robust indexing in difficult cases with crowded diffraction patterns (§6), explore difficult cases involving unit cell non-isomorphism (§7), and examine methods for handling indexing ambiguities in serial data (§8). Finally, we mention that our software suite can be applied to small-molecule diffraction (§9) in addition to macromolecules.

Importantly, this paper is not meant as a manual for *cctbx.xfel*. We have previously documented the software, including the *cctbx.xfel* Graphical User Interface (GUI) which allows for automated data processing of both small and large serial datasets using computing clusters (Brewster *et al*., 2016; Brewster *et al*., 2019*b*), and all parameters have detailed help messages accessible from the command line ^1^. Further, a new manual is being developed that will provide extensive documentation, examples, and use cases for the five existing hard XFEL sources and synchrotron use. Instead, this paper is meant to describe the algorithms and specializations needed to treat still images, as provided by our software.

## 2. Processing individual images with *dials.stills process*

The DIALS package can either be used as a series of standalone programs or as a library on which to build other programs and pipelines. For example, xia2 is an expert system for processing rotation (and more recently serial) diffraction data and can use either DIALS or XDS as its underlying pipeline (Winter, 2010). In our case, we created the wrapper program *dials.stills process* to read images (§2.1), and for each image execute a sequence of routines for spotfinding (§2.2), indexing (§2.3), refinement (§2.4), and integration (§2.5), using DIALS libraries. The program overrides library-default parameters to these algorithms, to enable stills-specific computation, as described below. Further, it provides extensive multi-processing capabilities to enable running in real time, even with fast data rates (§5).

For each image, *dials.stills process* normally runs the steps of import, spotfind-ing, indexing, refinement, and integration, but the exact workflow can be speci-fied with the `dispatch` parameters. For example, if initial indexing is being performed prior to refining detector geometry, with no need for Bragg spot intensities, `dispatch.integration=False` may be used to disable integration.

### 2.1. Data import

DIALS relies on accurate metadata in order to map pixel positions to laboratory space in three dimensions, and then to reciprocal space coordinates. This metadata includes detector position, pixel size, layout, sensor thickness and material, as well as beam direction and wavelength. DIALS uses the *dxtbx* library to import images, read their metadata, and load pixel values (Parkhurst *et al*., 2014). If the metadata isn’t accurate, for example if the detector position is wrong or has been refined separately, alternate geometry can be supplied, either using the `geometry or reference geometry` parameters.

Because import can be time consuming depending on hard drive and network speeds, care is taken so that processes in a multi-processing job (called “ranks” hereafter, using the Message Passing Interface (MPI) term for simultaneous processes) do not open the same image twice, nor read the same metadata twice. The image instead is held in memory during all steps of processing from import through to integration, then unloaded as the next image is loaded.

### 2.2. Spotfinding

Spotfinding in DIALS has been previously described (Winter *et al*., 2018) and is based on XDS (Kabsch, 2010). Generally, the same principles for identifying good spotfinding parameters for rotation series apply to stills, except that the detectors are often different. Rotation datasets at synchrotrons are usually taken using counting detectors with low background. Serial data from XFELs cannot use counting detectors due to the femtosecond scale duration of the pulses, therefore integrating detectors are used. In both cases, a critical parameter is detector gain. If the metadata is properly recorded, gain will be read from the image files. Often however it needs to be manually specified in two places, `spotfinder.threshold.dispersion.gain` and `integration.summation.detector gain`, the latter of which acts as a multiplier for variance in accordance with (Leslie, 1999). Gain can often be estimated with the program *dials.estimate gain*.

For integrating detectors, it is often useful to specify `spotfinder.threshold.dispersion.global_threshold`, which specifies a minimum value for pixels to be counted. This can help with separating background from dead pixels.

Most important however is the trusted pixel mask. In addition to the trusted range of the detector, an inclusive range within which pixel values are valid, a trusted pixel mask will eliminate unresponsive, noisy, or otherwise miscalibrated pixels. Several options are available to create them, including *dials.generate mask*, which can also combine multiple masks together, and the masking tool in *dials.image viewer*. In the case of no mask provided by the facility, a simple Python script can generate one from the standard deviation image generated by *dxtbx.image average*. In this example, only pixels with a standard deviation of less than 1 are marked as valid, which masks out noisy pixels with a high standard deviation:

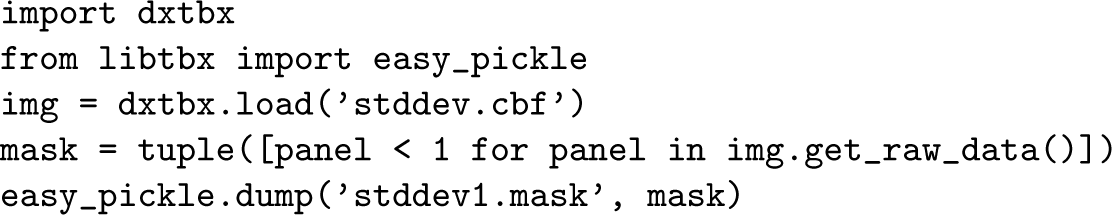

Hitfinding parameters can also be specified to ignore images with too few reflections, which can improve processing time. Hitfinding is controlled with the parameters under `dispatch.hitfinder`, and the default is to require at least 16 spots to index an image.

### 2.3. Indexing

Indexing is where the algorithms for stills begin to differ substantially from those for rotation datasets. Both *dials.index* and *dials.stills process* will auto-matically switch between the appropriate methods for still images by recognizing the lack of an oscillation angle in the image metadata. This behavior can be overridden with the parameter `indexing.stills.indexer`, chang-ing the default of `Auto` to either `stills` or `sequences`. It is gener-ally recommended to provide both `indexing.known_symmetry.unit` cell and `indexing.known_symmetry.space_group`, if known. This will greatly increase the indexing success rate (Hattne *et al*., 2014). If these are unknown, then after indexing in P1, the results can be clustered to find the best candidate cell and crystal system (Zeldin *et al*., 2015).

The stills indexer follows the processing procedure below, but first note that candidate crystal lattices are refined several times during the procedure. Refinement here means that the crystal lattice, both the orientation and unit cell, is optimized according to (Waterman *et al*., 2016), where the difference between the observed spot position and predicted spot position for each Miller index is minimized. The detector position is fixed in place, as is the beam model (direction and wavelength), as one image typically does not contain sufficient information to restrain these parameters (Brewster *et al*., 2018). Also, by default the cell is refined in *P* 1. This can be changed with `stills.refine_candidates_with_known_symmetry=True`, which will first apply space group symmetry prior to refinement.

The indexing steps are:

1. Attempt to find new lattices using available methods. This will produce a list of candidates, from which the algorithm will select the best given the criteria below. At present, two methods are available for indexing without knowledge of the crystal’s unit cell or space group:

- fft1d: the default for stills, this uses a hemisphere search for basis vectors by mapping spots on to candidate directions, and Fourier transforming them, looking for peaks that represent periodicity (Steller *et al*., 1997; Sauter *et al*., 2004).
- fft3d: performs a Fourier transform of reciprocal space directly (Bricogne, 1986; Otwinowski & Minor, 1997; Campbell, 1998; Otwinowski *et al*., 2012).

If the unit cell and space group are known, four additional indexing options are supported:

- real space grid search: given knowledge of the unit cell and spacegroup, this searches for a possible orientation to fit the data (Gildea *et al*., 2014).
- small cell: used for indexing still images from small molecule serial crystallography from as few as three spots (Brewster *et al*., 2014; Schriber *et al*., 2022).^2^
- low res spot match: matches low resolution spots to candidate indices. Designed primarily for electron diffraction still images.
- pink indexer: maps spot locations into trajectories of possible orientations that when intersecting, yield consistent crystal orientations (Gevorkov *et al*., 2020). It supports mono and polychromatic beams. Multiple methods can be attempted in sequence using the parameter *indexing.stills.method list*.
2. Assign Miller indices to spotfinder spots given the unit cell and orientation matrices found.
3. For each candidate, identify outliers where the spot prediction is too far from its observation, using Sauter & Poon (2010). This includes a preliminary round of refinement prior to outlier rejection, though these refined models aren’t retained here.
4. Refine each candidate and repeat outlier rejection.
5. Compute the two components of mosaicity used for still images, the average domain size and the distribution width of the relative orientations of the domains to each other, using Sauter *et al*. (2014). This includes outlier rejections for spots outside of the envelope in reciprocal space around the Ewald sphere predicted by the mosaic estimates.
6. Refine again, since outlier rejection will have affected the crystal parameters by removing outliers from the refinement target function. Then, recompute mosaic estimates using these best models. No further outlier rejection is applied.
7. Select the candidate with the best root mean squared deviation (RMSD) of the spots from their predictions to their observations, up to a maximum of 2 pixels.^3^
8. As a final step on the best candidate, repeat steps 3-6 to try and get any further improvement.
9. Try to find a new candidate by repeating the above steps if `indexing.multiple_lattice_search.max_lattices` is *>* 1. Any unindexed reflections will be used to attempt to find additional lattices from multiple crystal hits in the same image (Gildea *et al*., 2014).

Note, sometimes indexing with lower resolution data can give better results, using `indexing.refinement_protocol.d_min_start`. Note that the stills indexer doesn’t follow the pattern the rotation indexer does of multiple rounds of refinement with increasing resolution, so this parameter is only used to govern the resolution cutoff for indexing. Also note it is ignored for integration.

The default outlier rejection mechanism (Sauter & Poon, 2010) assumes an isotropic distribution of predictions away from reflections, but this is not the case for XFEL still images, where the wider bandpass of the XFEL source produces a distribution that has more displacement in the radial direction away from the beam than the transverse direction. This can be corrected using the mcd algorithm, and setting *positional coordinates* = *deltatt_t_ransverse* and *rotational coordinates* = *delpsidstar*. The effect on data processing is currently under investigation.

Finally, an additional indexing option is available, using known orientations. Here, a folder is specified with previous indexing results, and each image is matched up and assigned the indexing solution from the previous processing. This is useful in cases of multiple detectors, where one is used to get the indexing solution and then applied to another detector. See Figure 5 in Brewster *et al*. (2018).

### 2.4. Refinement

Even though refinement is done for most cases during indexing as noted above, an extra refinement step is provided, but disabled by default. It can be re-enabled using `dispatch.refine=True` (default is `False`) for cases such as using known indexing results or if refinement is disabled during indexing by setting `refine all candidates=False` (default is `True`).

### 2.5. Integration

Integration consists of three steps, all customized for still images:

- Profile fitting is disabled for still images, but even so the shape of the reflections is examined to determine the radial extent of the spots. Integration shoeboxes are sized accordingly.
- A list of Miller indices is generated to predict based on their proximity to the Ewald sphere, according to the mosaic parameters described above and in Sauter *et al*. (2014). Then, spot centroids are predicted by rotating them onto the Ewald sphere through the minimal rotation angle Δ*ψ*.
- Integration is performed using simple summation instead of profile fitting, with background subtraction done using a simple 2D plane fit as described in Leslie (1999).

As mentioned above, if the detector gain is not read from the metadata, it needs to be specified here as *integration.summation.detector gain*.

## 3. Time-dependent ensemble refinement

In Brewster *et al*. (2018) we showed that XFEL experiments tend to drift on the 10-100 *µ*m scale during long data collection, over the course of minutes, hours, or days. This can occur due to the sample interaction region moving during data collection (such as changes in the liquid jet direction), to sample reequilibration, temperature changes, or even settling in the experiment through a day/night cycle. These effects can be measured by batching the data as a function of time and refining the experimental models such as the detector distance for each batch. In each refinement, 100s to 1000s of crystals are simultaneously refined against a single detector model, such that each crystal informs the position of the detector, and the detector informs the unit cell dimensions and orientation of the crystal. Integrating with the refined detec-tor position can improve integration quality. We call this procedure time-dependent ensemble refinement (TDER).

TDER is available through the standalone program *cctbx.xfel.stripe experiment*, but typically it is performed as part of pipelines in the XFEL GUI, such that it occurs automatically during data collection.

## 4. Merging with *cctbx.xfel.merge*

*cctbx.xfel.merge* is the merging program for serial crystallographic data in *cctbx.xfel*,^4^ and it is modular in its design. It uses a series of worker modules to process data, with each worker executing a different task to, modify, filter, or merge the data. The workers are ran sequentially to read the data, scale it, and merge it, while computing relevant statistics, and they can be mixed, matched, and re-ordered depending on the specific use case. The default set of workers is:

- input: read data from disk in DIALS format.
- balance: balance input load. This is needed if the number of input files, which can contain integrated data from many images, is fewer than the number of MPI ranks being used.
- model scaling: build the full list of possible Miller indices and reference intensities, and set up the resolution binner for scaling and postrefinement.
- modify: apply polarization correction, re-indexing operators, and resolve any indexing ambiguities using libraries from *dials.cosym* (Brehm & Diederichs, 2014; Gildea & Winter, 2018).
- filter: reject whole lattices or individual reflections by resolution, unit cell, etc. The significance filter is recommended, which applies a per-image resolution cut-off estimated by where *I/σ_I_* falls below a given cutoff (0.5 by default, but often extended to 0.1).^5^ The unit cell filter is also recommended. It can be expressed either in terms of an absolute or percentage based cutoff away from unit cell parameter targets, or as a number of standard deviations (Mahalanobis distance) away from a multivariate Gaussian model of the unit cell target, derived from clustering the unit cells of all lattices with an algorithm like DBSCAN (Ester *et al*., 1996). For clustering, the closeness between two sets of unit cell parameters is evaluated with the NCDIST metric (Andrews & Bernstein, 2014).
- scale: scale the data to a reference dataset according to Hattne *et al*. (2014).
- postrefine: apply postrefinement, critically applying partiality corrections, scale factors, and B-factors (Sauter, 2015).
- statistics unitcell: unit cell averaging and statistics.
- statistics beam: wavelength averaging and statistics.
- model statistics: build full Miller list, model intensities, and resolution binner -for statistics. Can use average unit cell or a reference dataset.
- statistics resolution: calculate resolution statistics per crystal.
- group: re-distribute reflections over the ranks, so that all measurements of every HKL are gathered in the same rank, prior to merging.
- errors merge: perform error calibration according to one of three possible methods, both described in Brewster *et al*. (2019*a*):

– `Ev11`: adjust individual measurement errors using three terms: *s_fac_*, *s_B_*, and *s_add_*, refined according to Evans (2011) until the errors better explain the observed variance in the data. Merged intensities will then be a weighted mean using inverse variances as the weights.
– `MM24`, the default: further development of the `Ev11` model that differs in two primary ways: 1) *s_add_* is parameterized to assign varying levels of error to each lattice using its correlation to the supplied reference dataset, and 2) the target function is reformulated to increase optimization robustness (Mittan-Moreau *et al*., 2025).
– `errors_from_sample_residuals`: variances for merged data are the variance of the unmerged observations for each HKL. Merged intensities are then an unweighted mean of the unmerged observations, as originally proposed in Schwarzenbach *et al*. (1989).
- statistics intensity: calculate resolution statistics for intensities.
- merge: merge HKL intensities and output “odd”, “even” and “all” HKLs as mtz files. The user may decide to either merge the Friedel mates or not, depending on if anomalous differences are important to retain.
- statistics intensity cxi: compute *CC*_1_*_/_*_2_ and related statistics.

The list of workers can be changed with the parameter `dispatch.step_list`.

### 4.1. Merging use cases

#### 4.1.1. Scaling and merging separately

The *cctbx.xfel* GUI (Brewster *et al*., 2019*b*) scales and postrefines each set of data separately prior to merging all data together, since at this stage images are still independent, as scaling and postrefinement depend only on the image itself and the reference dataset. This increases efficiency as it allows for each image to be only scaled and postrefined once. As the dataset grows during data collection, the merging can be done repeatedly, and the user can monitor improvement in data quality.

This is configured at the ‘run’ level, where a run is usually around 5-10 minutes of continuous data collection. Runs are scaled and postrefined, using the workers listed above: starting at input through postrefinement plus beam, model and resolution statistics. Then the merging program is utilized again, this time with all the runs to be included in the dataset, using the input and model scaling workers to load the data and reference structure, then skipping to the group, errors merge, and merge workers, including any dataset as a whole statistics workers. Merging in this way ensures that as new data are collected, data that has already been scaled and postrefined is not scaled and postrefined again.

#### 4.1.2. Merging without a reference

In the case of not having a scaling/merging reference dataset, a ‘bootstrap’ procedure can be used, similar to the one used by PRIME (Uervirojnangkoorn *et al*., 2015) or in Brewster *et al*. (2019*a*). First, use the ‘`mark1`’ scaling algorithm which performs a simple average of the data, and the list of workers does not include postrefinement. The parameter `scaling.model` is not specified, and as error model we use `errors_from_sample_residuals` instead of the default `Ev11`. This will create a new mtz file, which can then be used as a reference in subsequent merging using the default ‘`mark0`’ scaling algorithm, which scales the data to a reference dataset. We recommend using several iterations. For example, use a `mark1` merge followed by several `mark0` merges, each using the output mtz from the previous merge as a reference. *CC*_1_*_/_*_2_ should improve over the first few cycles and converge.

### 4.2. Merging statistics

*cctbx.xfel.merge* produces a log file named `main.log` with statistics about the merged dataset. Here we walk through the critical tables and values, indicating how to read important numbers and providing guidelines for estimating crystal quality and resolution cutoffs.

#### 4.2.1. Unit cell statistics

The statistics unitcell worker will average all the unit cells in the dataset and produce a histogram of the unit cell lengths with a standard deviation. In the case of unit cell non-isomorphism, where filtering or clustering are important, these statistics can be useful to judge whether the merged dataset has isomorphous unit cells.

#### 4.2.2. Lattice resolution

The statistics resolution worker produces a table of the number of lattices accepted by the significance filter at each resolution bin and the percentage that represents of the total number of lattices. A sample with a high percentage of accepted lattices at high resolution is a high quality sample, and fewer lattices will be required for a complete dataset. A sample with a lower percentage of accepted lattices may still yield a high quality dataset, but it’s likely more images will need to be merged to bring up the multiplicity, which can add noise to the final data.

#### 4.2.3. Intensity resolution

The statistics intensity worker produces three tables, two for odd and even numbered lattices, one for all lattices. These tables show by resolution the completeness, multiplicity, number of measurements, and *I/σ_I_* of the dataset. Because of the significance filter, multiplicity will represent the number of measurements accepted at a given resolution bin; see the section on Data quality evaluation, including figures S2 and S3 in Ibrahim *et al*. (2020).

One of two rules of thumb for resolution cutoff is in this table: the resolution where multiplicity falls below 10×. This rule of thumb applies when using the significance filter, and it comes from experience in merging many datasets, and not from a rigorous statistical model. As always, good judgement should be used when choosing a resolution cutoff, taking the statistics of a dataset as a whole into account and judging significance by map quality. The gold standard of resolution estimation remains the paired refinement procedure of Karplus & Diederichs (2012). Note also that *I/σ_I_* is not used as a cutoff, due to often weak data containing useful signal and the difficulty in modeling error in serial crystallography (Brewster *et al*., 2019*a*).

#### 4.2.4. Merging statistics

The last worker, statistics intensity cxi, produces the final table which includes *CC*_1_*_/_*_2_ and Rsplit. The second rule of thumb for resolution cutoffs is applied here using *CC*_1_*_/_*_2_, but rather than a specific cutoff value, we generally cut the data off at the resolution where *CC*_1_*_/_*_2_ ceases to decrease monotonically, even though that can sometimes reach as low as 0%. The 10× multiplicity cutoff is generally lower resolution, but *CC*_1_*_/_*_2_ remains a critical measure, especially the overall value which should be greater than 95% in most cases.

Our two rules of thumb, 10× multiplicity and monotonic *CC*_1_*_/_*_2_, are useful because during beamtime where fast feedback is needed, these figures can be determined rapidly. However, we wanted to investigate the correlation between them and paired refinement results, so we performed paired refinement on 3 XFEL datasets (see section 11). We found that the paired refinement results agreed with the 10× multiplicity estimate to within +/-one resolution bin, and therefore it seems like a useful heuristic.

Notably, *CC*_1_*_/_*_2_ is very sensitive to outliers, even from single reflections. Oversaturated spots or badly masked, noisy pixels can greatly affect *CC*_1_*_/_*_2_ in certain cases. If *CC*_1_*_/_*_2_ dips or is otherwise suspicious for a single bin, or if the overall *CC*_1_*_/_*_2_ seems too low for the number of images merged, removing data until the culprit is found (could be a single image or even a single reflection) is critical. Approaches for finding these outliers have been described (Assmann *et al*., 2016; Assmann *et al*., 2020), and are implemented in *cctbx.xfel.merge* in the optional *deltaccint* worker. This worker uses the *σ* − *τ* method for determining *CC*_1_*_/_*_2_, which does not rely on splitting the dataset into random halves, to compute a change in *CC*_1_*_/_*_2_ when leaving out each lattice one at a time. An example from a Photosystem II dataset is shown in Figure 1, where a single image that causes a dip at 2.2Åneeds to be removed. The *deltaccint* worker computes the *CC*_1_*_/_*_2_ values when removing each lattice in the dataset one at a time, then reports worst 30 images in terms of their contribution to *CC*_1_*_/_*_2_ and identifies an inter-quartile range (IQR, or the third quartile (q3) minus the first quartile (q1)) mul-tiplier that can be used to remove them. In this case a multiplier of 100 removes the problem image, meaning all images with a Δ*CC*_1_*_/_*_2_ *> q*3 + *IQR* ∗ 12 are removed. 100 was determined by running the program once and examining the log, then re-running with the parameter *statistics.deltaccint.iqr ratio* = 100. Note that *CC*_1_*_/_*_2_ value is the overall value for all the data, rather than the average the value of each resolution bin, as proposed in Assmann *et al*. (2016). Note also that the new error model, MM24 (see error merge above), can also help in situations like these, as it can down-weight the affected image using its correlation to the supplied reference dataset (Mittan-Moreau *et al*., 2025).

**Fig. 1.**
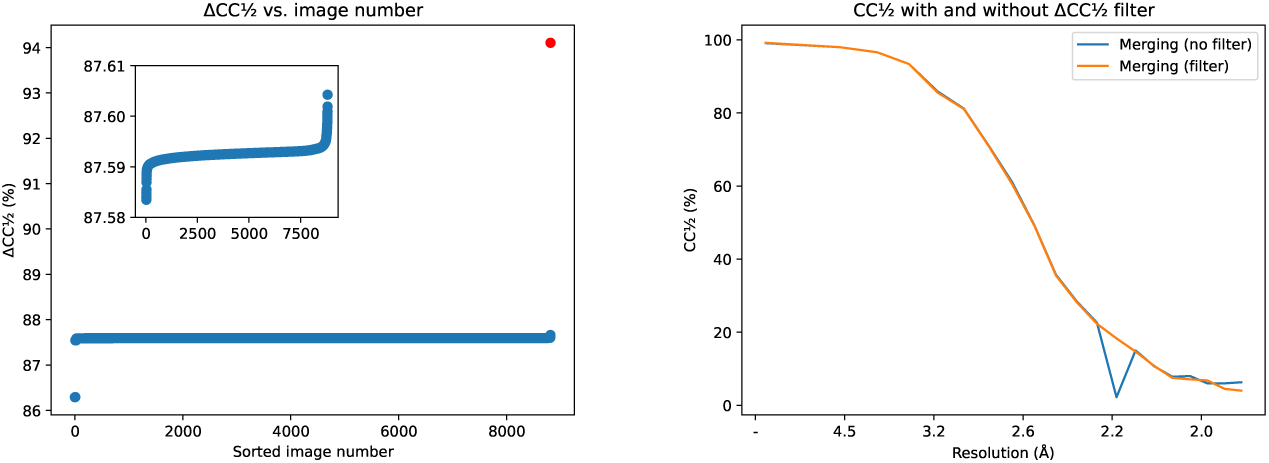
Left: sorted *CC*_1_*_/_*_2_ for an 8816 lattices Photosystem II dataset. A critical outlier is labeled with a red dot. Inset: tight zoom on the Y-axis. Right: *CC*_1_*_/_*_2_ vs. resolution with and without the outlier

Importantly, once a resolution cutoff is determined, data should be re-merged using this cutoff instead of applying a cutoff to the data in down-stream steps, such as protein structure refinement. This is to remove poor data from the merging process where it can affect procedures such as scaling and postrefinement.

## 5. Multiprocessing and the XFEL GUI

In addition to processing images, *dials.stills process* further aggregates and marshals data processing using a variety of multiprocessing methods, including local Python multiprocessing on a single node, and processing on multiple nodes using MPI. As the images are independent of each other, images are fully processed one at a time. Importantly, not all images will successfully pass each processing step, *e.g.* in most serial datasets not all images have crystals and so some images will fail to index. This has implications on how data are divided in multiprocessing. For all multiprocessing modes except MPI, data is “striped”, meaning the images are evenly divided between all processes using a round-robin method. This accounts for the common pattern of sections of “hits” and “misses” in the data. An even distribution over time would mean some processes would finish early, but a round-robin striping will evenly distribute hot-spots of crystals in the data.

However, even in striping it is common for some processes to finish early, so in MPI mode, *dials.stills process* instead dedicates one rank as a server and the other ranks as clients. The server distributes images to each of the clients, which process the images and report back to the server for more. This ensures all ranks are busy until the job is fully processed. We have used this approach to process data at speeds up to 9 kHz to provide live feedback for many XFEL experiments (Blaschke *et al*., 2024).

MPI has also been applied to *cctbx.xfel.merge* to merge huge datasets on supercomputers, including for example a photosystem II dataset with over 600,000 images and over 10^9^ measurements.

The *cctbx.xfel* GUI is designed to allow configuration of processing pipelines and has been used at each of the current five XFELs and at synchrotrons. It has been recently updated to be more user-friendly and intuitive, with tooltips and documentation to allow configuring the most important and common parameters while allowing advanced processing techniques for difficult cases. Generally we have found there are two main challenges when using the GUI during and after experiments: calibration and batch processing using a supercomputer.

### 5.1. Calibration

Calibrating XFEL detectors is challenging, facility specific, and generally beyond the scope of this paper. Brewster *et al*. (2018) provides detailed examples on how to refine the geometry of a CSPAD detector and is generally useful for understanding the methods involved. However, the upcoming *cctbx.xfel* manual contains details specific to the current known instruments, including reading XTC-format filestreams using the psana library at LCLS for the CSPAD, Rayonix, Jungfrau, and ePix detectors, reading SACLA Rayonix and MPCCD HDF5-format data, creating NeXus files (Bernstein *et al*., 2020) for PAL Rayonix and Jungfrau data, SwissFEL Jungfrau data, and EuXFEL AGIPD and LPD detector data, and calibrating wavelengths using X-ray spectrometer data from *e.g.* LCLS and SwissFEL, including pedestal correction.

### 5.2. Computer cluster support

X-ray facilities vary in how they provide computing support. LCLS recently introduced the Stanford shared data facility (S3DF), or data can be live transferred to NERSC for processing using reserved resources (Blaschke *et al*., 2024). Both approaches can yield live feedback, but NERSC is scalable to fast data rates beyond 120 Hz and may be necessary for more intensive tasks such as random sub-sampling (see below). SACLA has a small cluster available on site, as does SwissFEL. PAL has access to Kisti, and the EuXFEL has their Maxwell cluster.

Each of these computing environments has unique demands for submitting and running jobs using multiple nodes. There are different job submission environments such as slurm, PBS, and HTCondor, and different requirements for building *cctbx* and using MPI. There are different approaches for monitoring for data and different sets of configuration parameters needed to read data. The *cctbx.xfel* GUI supports being configured for all of these systems using simple interfaces, and once configured, the rest of data processing is standardized. Please see the *cctbx.xfel* manual for details.

## 6. Indexing crowded patterns with random sub-sampling

To demonstrate the flexibility of *cctbx.xfel* we present a new method, random sub-sampling, which we have been using to increase the indexing success rates for difficult datasets. High resolution, high-quality images taken on single panel, centered detectors positioned to capture both low and high resolution data are generally trivial to index for most cases. However, there are many situations where these assumptions are broken. 1) Often crystal concentration is high enough to produce more than one or two lattices per image. In the case of large unit cells, this can generate thousands of spots on an image and defeat Fourier methods. 2) Certain experiments use complex geometries, including offset and off-center detector positions where only a portion of the pattern is collected. Often not enough reflections are captured to index a lattice. 3) Smaller unit cells, on the order of between 30-50Å on a side, can generate sparser patterns. Not so sparse that entirely different methods are needed (see §9), but such that only 20-50 reflections are recorded. This also can lead to not enough reflections to index the image.

Random sub-sampling operates under the assumption that using all the available reflections can lead to either too much ambiguity in Fourier space or random lattice poisoning in the case of too few spots. However, if false or extraneous signal can be removed, indexing can succeed. It is difficult to know which sets of reflections must be removed. Random sub-sampling removes a percentage of the spotfinder spots at random if indexing fails and attempts indexing again. The default protocol is enabled with `indexing.stills.random_sampling.enable=True`, and it operates by removing 2% of the data at a time, down to 50%, for a total of up to 25 indexing attempts per image (these parameters are customizable). If at any point indexing succeeds, the procedure stops and the program moves on to the next step (refinement or integration). Importantly, at each step the full set of strong reflections is sampled, allowing discovery of the best set of reflections as the dataset size is restricted.

After applying sub-sampling, almost 44% of the images can be indexed, increasing the number of indexed images by 34% over the defaults. Importantly, this includes some of the images with over 1500 spots that couldn’t be indexed previously. The real space grid search (RSGS) algorithm has been highly successful for rotation crystallography in finding multiple lattices from datasets where up to 6 crystals were exposed simultaneously (Gildea *et al*., 2014). We have used it previously in similar situations in serial crystallography, and wanted to compare its performance for this use case. Immediately we see it indexes more images that FFT1D (Figure 2, third row), and over the whole run it increases to 47%. With sub-sampling 57% of the images are indexed.

**Fig. 2.**
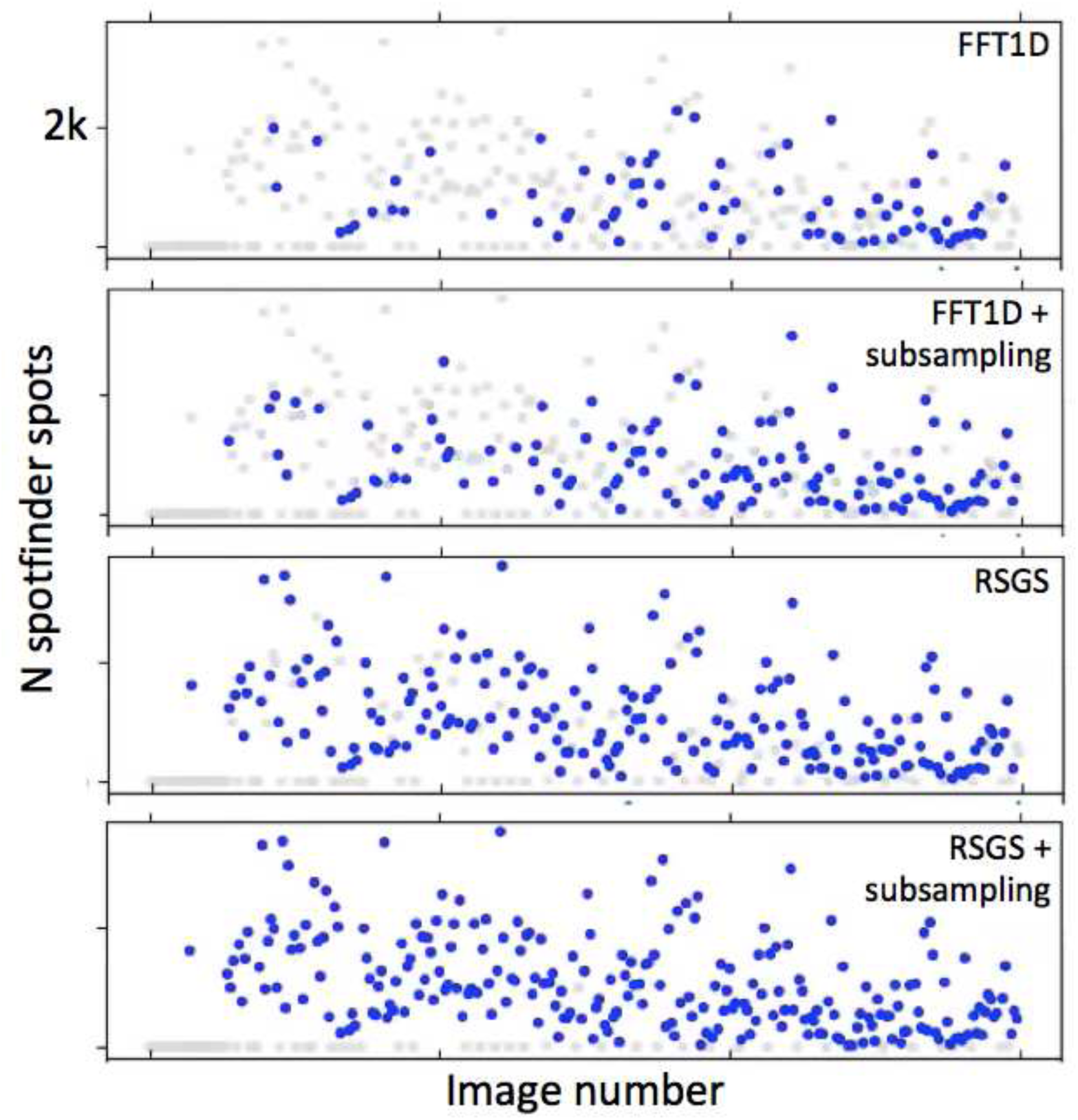
Indexing plots from the *cctbx.xfel* GUI. Four indexing trials are shown. One dot is shown for each of about 300 of the 10536 images from run 62 of LY64, plotted as the number of spotfinder spots vs. image number. Blue dots: indexed images. Gray dots: images that failed to index.

We investigated these methods further by checking merging statistics including the anomalous peak height for each combination of these methods, plus 4 more tests, where we tried both FFT1D and RSGS for each indexing attempt (this can be enabled using the `indexing.stills.method_list` parameter) (Table 1 and Figure 3). We found that the highest anomalous peak heights and best merging statistics were achieved using combinations of indexing methods and sub-sampling. We encourage users to experiment with these parameters to best fit their data, and for large unit cells, depending on computing available, to even consider greatly increasing depth of sub-sampling. For example, we have seen for very large unit cells (*>*300Å), that a sub-sampling regime of 100% down to 20% with 1% steps, and three random attempts per step, can increase indexing success, even though this yields 240 attempts per image.

**Fig. 3.**
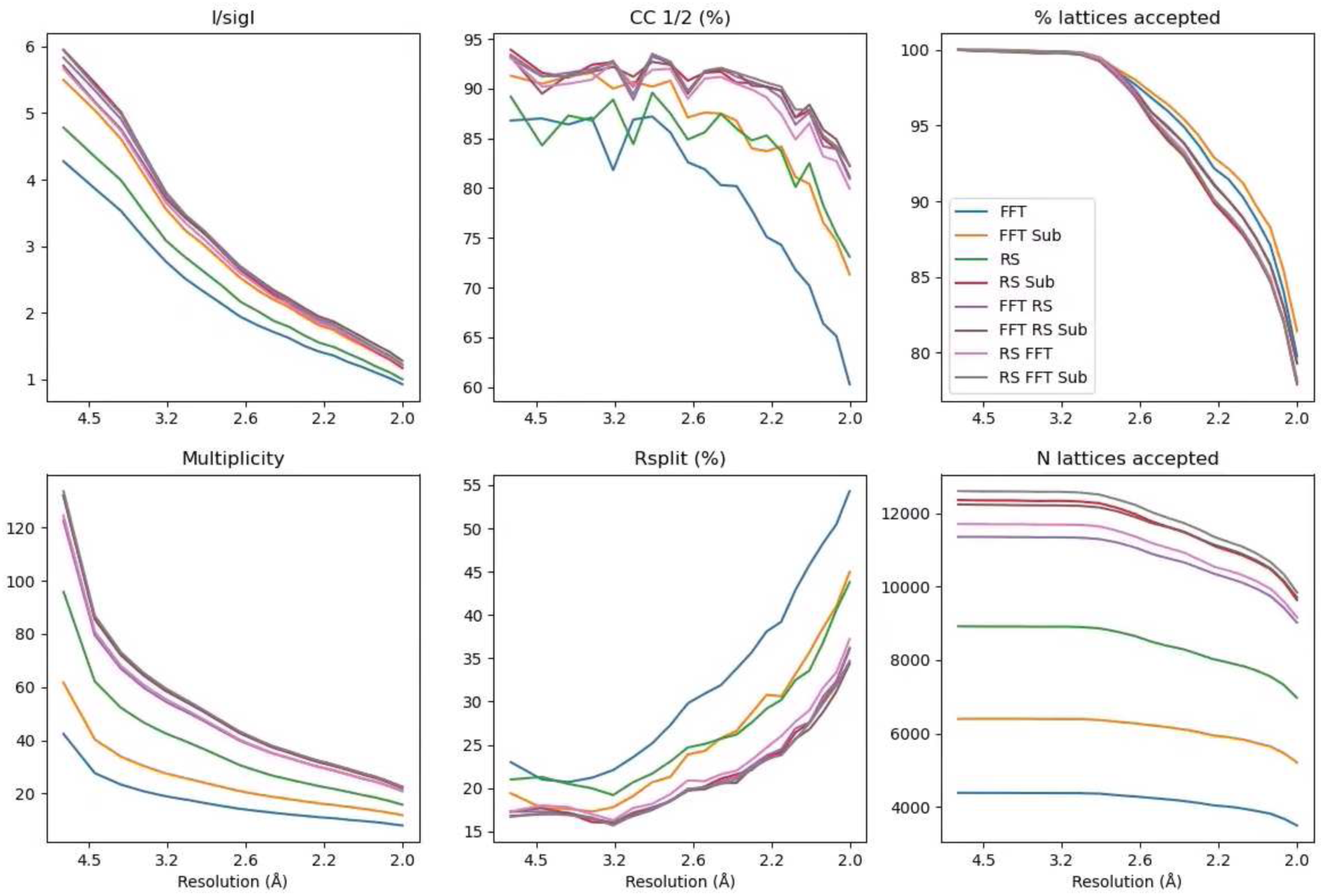
*I/σ_I_*, *CC*_1_*_/_*_2_, % lattices accepted by the significance filter, multiplicity, Rsplit, and N lattices accepted vs. resolution for each of the 8 MCR trials

**Table 1.** Random sub-sampling trials.

| Trial | Method(s) | Sub-sampling | N indexed | N lattices | Ni 1* | Ni 2* |
| --- | --- | --- | --- | --- | --- | --- |
| 1 | FFT1D | No | 5617 | 8122 | 10.54 | 9.08 |
| 2 | FFT1D | Yes | 7988 | 11330 | 12.62 | 11.47 |
| 3 | RSGS | No | 10061 | 18094 | 11.34 | 9.43 |
| 4 | RSGS | Yes | 12264 | 21149 | 12.29 | 9.27 |
| 5 | FFT1D + RSGS | No | 10694 | 17720 | 12.37 | 10.16 |
| 6 | FFT1D + RSGS | Yes | 12314 | 19891 | 12.61 | 10.31 |
| 7 | RSGS + FFT1D | No | 10694 | 18972 | 11.90 | 9.62 |
| 8 | RSGS + FFT1D | Yes | 12314 | 21173 | 11.82 | 9.30 |

\*Anomalous peak height for nickel atoms in MCR. For 7 and 8, the order of FFT1D and RSGS was swapped.

## 7. Unit cell non-isomorphism and clustering

Unit cells in serial crystallography are estimated from the periodicity of the lattice, usually using Fourier methods. Generally two of the axes are more orthogonal to the beam, but the third axis is generally parallel to the beam. This leads to fewer planes being measured in reciprocal space and a less well measured axis. This is illustrated in Figure 4 and is treated in Figure 6 of Brewster *et al*. (2018).

**Fig. 4.**
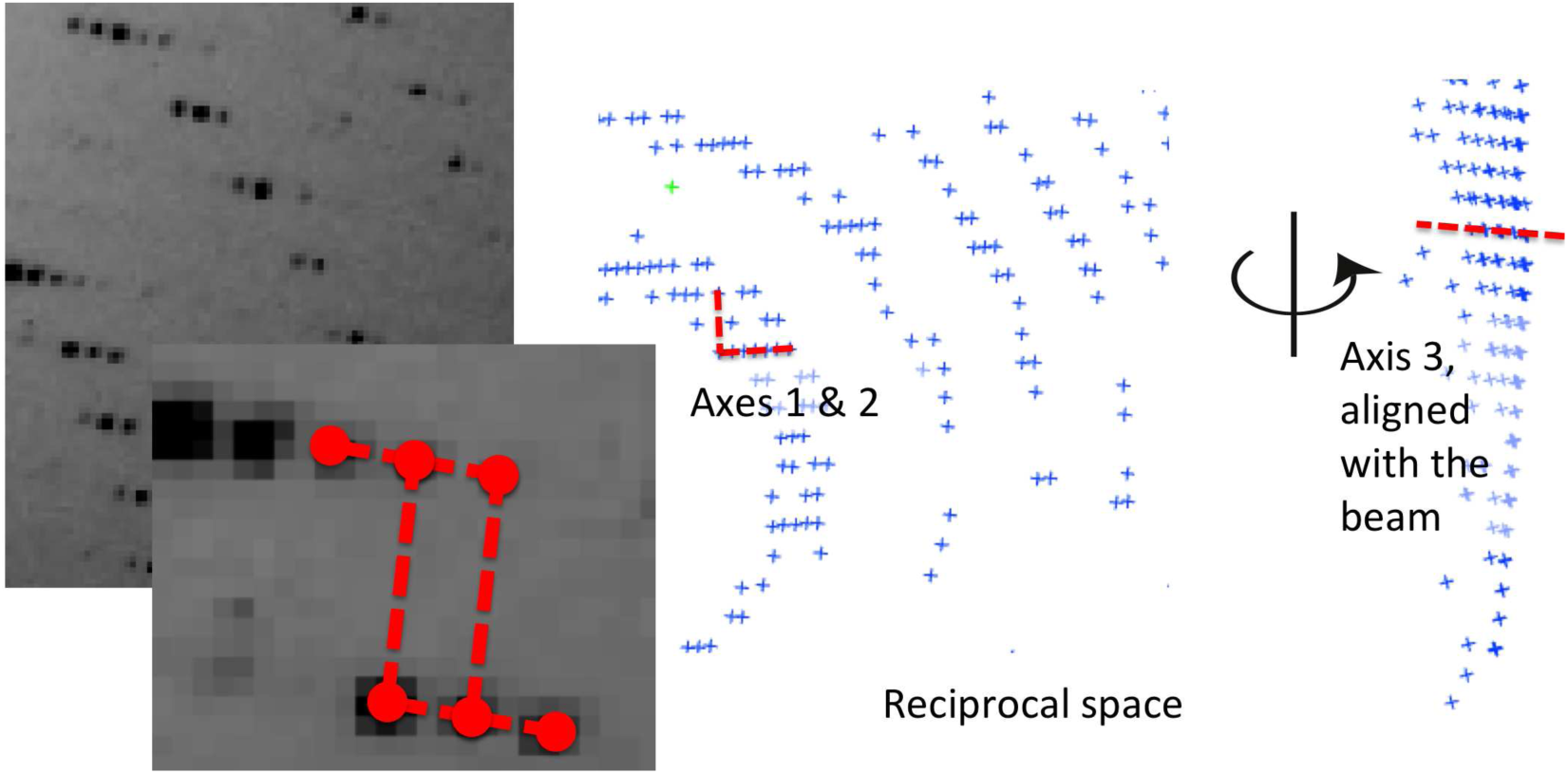
Stills have less 3D information. Left: photosystem II image with inset zoom. Red dots and lines are lattice directions. Right: spotfinder spots in reciprocal space, with a 90° rotation to visualize that lunes represent slices through the Ewald sphere.

**Fig. 5.**
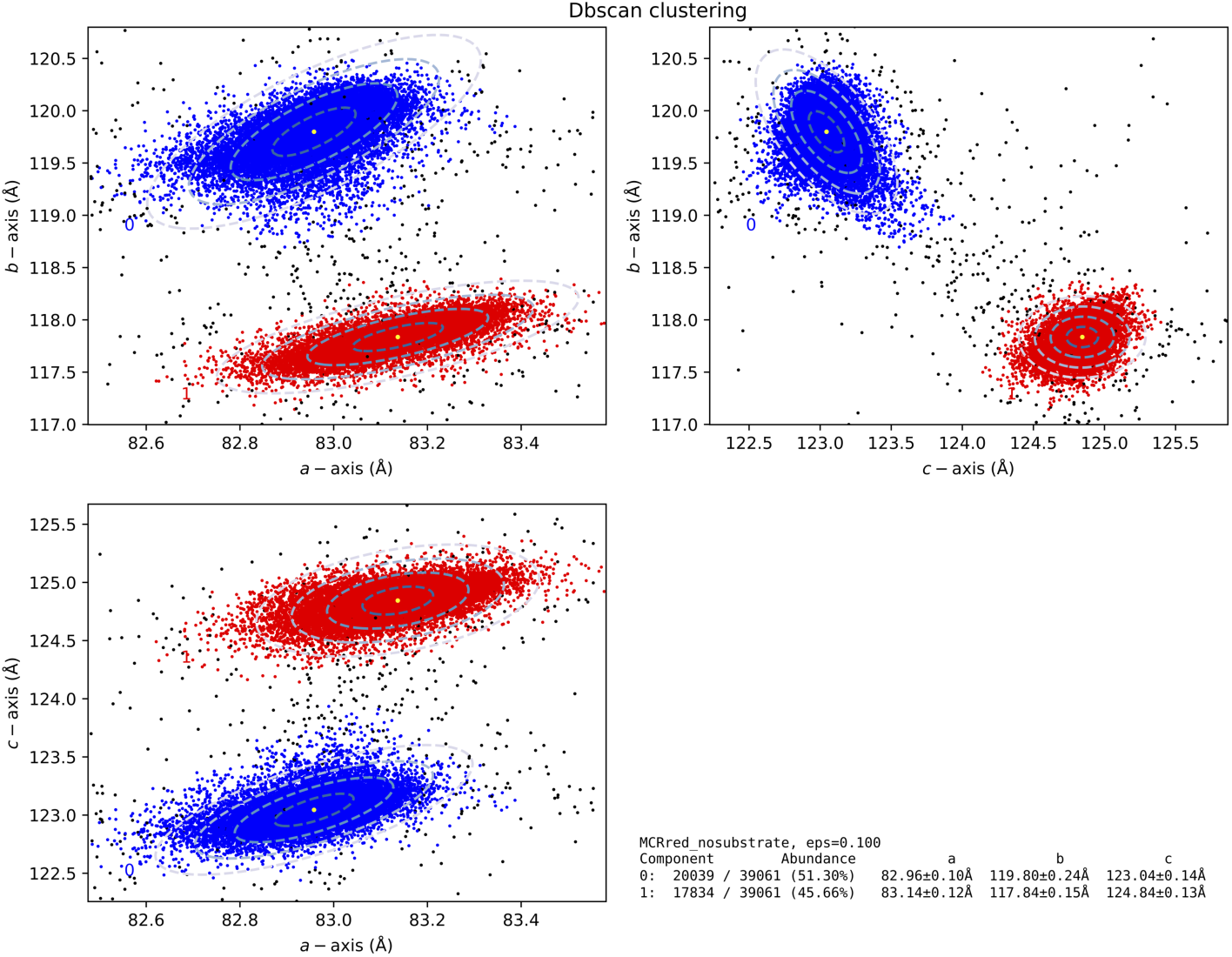
Unit cells from LCLS experiment LY64 as displayed by the *cctbx.xfel* program *uc metrics.dbscan* with an `epsilon`value of 0.1

**Fig. 6.**
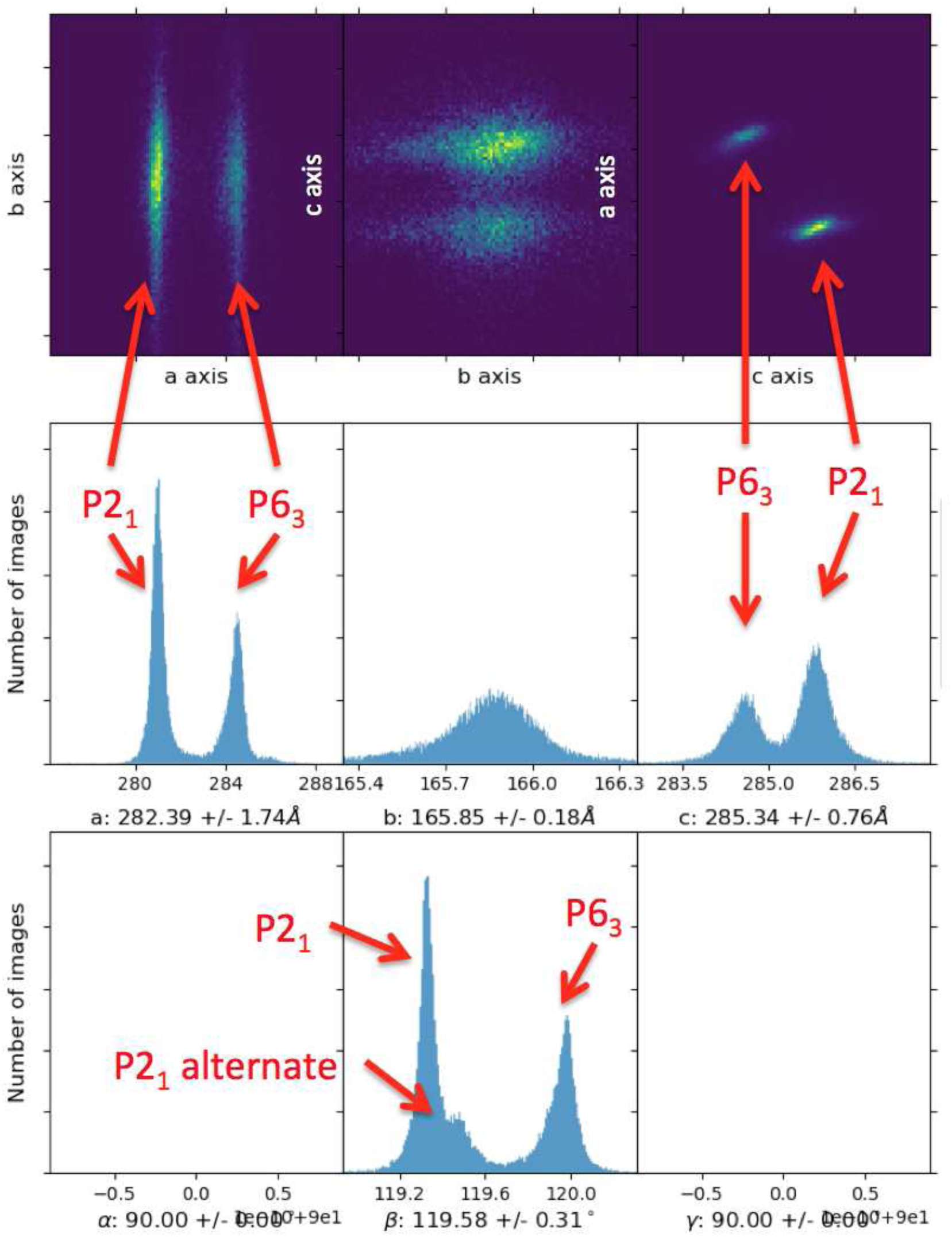
25,265 Photosystem I cells indexed in *P* 2_1_ and histogrammed using *cctbx.xfel*.

Very accurate detector geometry can correct for this (Brewster *et al*., 2018), but enough measurement error exists on all three axes that unit cells form distributions. When merging there are two ways to deal with this problem:

1. Use a unit cell length filter to remove any cells within a cutoff (percentage or absolute in Å). This creates a flat box of cells in a plot of *a* vs. *b* vs. *c* that does not follow the reality of most serial experiments, where radial error from inaccuracies in wavelength and detector distance contribute a uniform distortion of the cell dimensions, forming elliptical clouds of unit cell parameters oriented toward the origin.
2. DBSCAN clustering using multivariate ellipsoids. Currently, the features used for cluster classification are the unit cell parameters *a*, *b*, and *c*, and the relevant distance metric is the L2 norm.^6^ Polytope boundaries in unit cell space are not treated, as would be the case with a more complex metric. The “pros” to this are that we can use a standard clustering algorithm from scikit-learn. The “cons” are that we cannot support triclinic, monoclinic, or pseudo-symmetry. This is a matter of active research in our group. Current work includes using the Andrews-Bernstein metric NCDist which can help resolve polytope boundaries, and using it as a precomputed metric for DBSCAN from scikit-learn. We are also testing the Rodriguez-Laio method, including PCA and N-dimensional embedding for visualizing four-or six-feature clustering jobs for monoclinic or triclinic cells.

DBSCAN uses a parameter `epsilon`, which is the characteristic maximum distance above which items are no longer considered part of the same cluster. The idea here is that `epsilon` makes this an interactive program: users are asked to run the program with several different values of `epsilon` and then choose the clustering result that most closely matches with visual perception (Figure 5). Users should especially notice the cluster centers, as well as the standard deviation contours that should nicely follow the clusters. If unit cell instances are fairly sparse and/or no clusters are labeled, the `epsilon` parameter should be increased. If however, it appears that a labeled cluster should actually be two separate components (perhaps with unequal populations) the `epsilon` parameter should be decreased. The user should note the component number and sigma level that matches the cluster of interest. Then, during merging, this component can be separated from other lattices given the sigma level. The interactive program is available in the XFEL GUI or command line (see *cctbx.xfel* manual).

We especially encourage users with difficult cases including pseudo-symmetry in monoclinic and triclinic cells to contact the authors, and to make their data publicly available in repositories such as CXI.DB (Maia, 2012).

## 8. Indexing ambiguities

Serial datasets in Laue classes −3, −3m1, −31m, 4/m, 6/m, and m-3 are known to be intrinsically challenging because the lattice has a proper rotation (or two, for trigonal crystals) that is missing from the crystal point group. In single-crystal experiments with these polar space groups, this higher lattice symmetry causes the common phenomenon of merohedral twinning. In serial datasets, each individual frame may be indexed in two distinct orientations related by a merohedral reindexing operator. Since the lattice fits identically well in both possible orientations, the indexing ambiguity can only be resolved by examining the observed intensities.

Several approaches have been described for resolving ambiguous indexing in serial diffraction. Most simply, the intensities in a frame can be correlated against a reference structure in each possible orientation. The orientation that gives the better correlation to the reference structure is selected. This approach is not suitable for unknown structures, and is highly susceptible to weak and noisy data as are commonly collected in serial experiments. In 2014 an improved method was reported by Brehm and Diederichs in which correlations are measured between individual frames and the frames are subsequently embedded in a vector space with dimensions matching the order of the indexing ambiguity. Starting from random coordinates in the n-dimensional vector space, a minimization step is performed in which the dot products of vectors approximate the correlation coefficients between the corresponding frames. After this minimization, frames which share a common indexing sense are close together (i.e. highly correlated), and differently indexed frames are far apart (i.e. uncorrelated) (Brehm & Diederichs, 2014). This algorithm is structure-agnostic and is highly tolerant of noisy measurements (Diederichs, 2017).

Two algorithms are available in *cctbx.xfel.merge*:

1. `modify_reindex_to_reference` steps through each diffraction pattern individually, correlating the Bragg intensities with 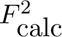 from a structural model. Clearly there are disadvantages: the data may not be exactly isomorphous to the reference, so there may be false assignments of the indexing sense. In fact, the data analyzed for the first use case below produced correct alignments only 85% of the time.
2. `modify_cosym` performs a mutual alignment of all the still shots, using an *N* × *N* matrix of correlation coefficients (*N* is the number of diffraction patterns). The approach was first proposed by Brehm and Diederichs to address the indexing ambiguity problem (Brehm & Diederichs, 2014). Later, Gildea and Winter expanded the concept to allow both the alignment of polar cells, as well as the *de novo* determination of Laue symmetry (Gildea & Winter, 2018).

Two use cases are presented here as examples, for treating experimental cases where there is an indexing ambiguity.

### 8.1. Use case 1: Simple polar group

First we describe the mutual alignment of an 8000-image dataset with space group *P* 6_3_, merged in about 30 minutes on a 60-core AMD server. Data files were previously indexed and integrated with *dials.stills process* in space group *P* 6_3_. The modify cosym worker is added before scaling, and if using `reindex_to_reference`, a reference struc-ture must be supplied to compare the intensities of each image to. If using cosym, the reference structure is unneeded, and instead the space group can be supplied if known. If the space group is unknown, it will attempt to be determined according to Gildea & Winter (2018), otherwise, the clustering algorithm in Brehm & Diederichs (2014) is used directly to align images to each other using intensity measurements. The two approaches can be directly compared using a script described in the manual.

If a reference structure is supplied, to avoid bias cosym will not use it for alignment, until the final step where one set of images needs to be reindexed to match the other set (in the case of a 2-fold ambiguity). The set with the closest CC to the reference is chosen as the anchor dataset and the other set is reindexed onto the anchor. This allows 1) mark0 merging, especially if postrefinement is used, since the data are aligned to the reference, and 2) anchoring the output against the reference allows the merged data to be dropped into the refinement program directly for isomorphous refinement, without a molecular replacement step.

In this example, the program is ran using 60 MPI ranks. There is a critical tradeoff involving the number of MPI ranks. Analysis of a large dataset (*N* = 8000) would be prohibitive if the full *N* × *N* matrix were to be analyzed. Instead, we break the data into *n* tranches of size *T* = *N/n* shots, giving *n* = 133. In practice each MPI rank is given 3 tranches to analyze, or 400 × 400 experiment matrix in this case, including overlap with 2 other ranks, so that the polarities determined within each rank can be mutually reconciled by a 3-way vote. In other words, the indexing polarity of every image is determined 3 times by including the image in three different tranches. The polarity is accepted if either all three tranches produce the same polarity for the image (consensus voting, the default), or if 2 out of 3 produce the same polarity for the image (majority voting). Useful tranche sizes *T* range from about 100 to 600. Smaller *T* produces drastically shorter wall clock time, while larger *T* produces dramatically superior Brehm-Diederichs embedding plots, as in Figure 4 of Brehm & Diederichs (2014). The embedding plot (see `modify.cosym.plot.interactive`) must be checked to ensure clusters are well separated from the 45-degree diagonal, and that the cluster centers are at a good distance from the origin (0.4-0.8 is good). If the clusters look bad, the tranche size should be increased. Note, the algorithm will not work unless *n* ≥ 5. Some additional parameters are described here:

- `modify.cosym.dimensions=2`: this is absolutely critical. We set this to 2 dimensions for space group *P* 6_3_, as there are exactly two groups (cosets) expected in the final sort. Setting this to the default (auto determine) has the consequence of performing the embedding analysis in a higher dimensional space (6) where the clusters cannot be found. Keep this at 2 for most cases, or 4 for space groups *P* 3, *P* 3_1_, and *P* 3_2_.
- `modify.cosym.min pairs=3`: minimum number of mutual Miller indices per-lattice and per-symmetry-operator (symop) to form a correlation-coefficient cross-term with another lattice/symop combination. Takes the default from Gildea & Winter (2018); increasing this will drastically reduce the data used for alignment, due to fewer pairs of images being able to be aligned, likely to the detriment of the final results.
- `modify.cosym.weights=count`: This is critical. Setting weights=None makes the cosym algorithm fail, while the other options have not been successfully tested for stills.
- `modify.cosym.plot.interactive=True`: this displays an embedding plot as in Brehm & Diederichs (2014) to assess whether the multiple indexing solutions are sufficiently resolved from each other. The embedding plot is a useful tool for setting the number of MPI ranks (and thus the tranche size for analysis). For routine work, this may be omitted, and a plot will be saved in the output directory.

### 8.2. Use case 2: P 6_3_/P 2_1_ mix

Photosystem I (PSI) crystallized in *P* 6_3_ was an original driver for the Brehm and Diederichs algorithm for resolving polar space groups in serial data (Chapman *et al*., 2011; Brehm & Diederichs, 2014). However, it has also been reported in *P* 2_1_ (Gisriel *et al*., 2019), and while our recent report showed PSI again crystallized in *P* 6_3_ (Keable *et al*., 2021), our group has unpublished data with a crystal condition in which both isoforms can be observed in the same sample. The two forms are closely related. A quick analysis with the program `labelit.check_pdb_symmetry` (Poon *et al*., 2010) from the PHENIX suite (Liebschner *et al*., 2019) shows that the monclinic *P* 2_1_ form can be morphed into the hexagonal *P* 6_3_ form with just a 1.3° lattice distortion and an r.m.s. *α*-carbon displacement of 2.2 Å.

Resolving the polar ambiguity in *P* 6_3_ was described above, but another complication arises when indexing datasets in which the higher metric symmetry is not exact (merohedral) but approximate (pseudomerohedral). A familiar example from chemi-cal crystallography is a monoclinic crystal with *β* coincidentally close to 90°. In this situation the Laue symmetry is 2/m and the lattice pseudosymmetry is mmm. In single-crystal diffraction, twinning may occur by rotation around a pseudo-twofold axis of the lattice, and in serial diffraction, indexing will be ambiguous via reindexing by the same rotation. Since the higher lattice symmetry is only approximate, there is often evidence of this phenomenon via splitting of peaks in a single-crystal pattern or splitting of refined unit-cell distributions in a serial dataset.

In monoclinic Photosystem I, the lattice is pseudo-hexagonal with *a*, *b*, *c* = 281, 166, 286 Å and *α*, *β*, *γ* = 90, 119.4, 90°. The Laue class of the structure is 2/m (order 4) and the lattice pseudosymmetry is 6/mmm (order 24); thus, in principle, six mutually inconsistent indexing senses could be observed.

In order to treat these data we first used the clustering algorithm described in section 7 to separate the hexagonal from the monoclinic lattices (Figure 6)^7^. Next, we expected that monoclinic Photosystem I might also display an indexing ambiguity since its structure and lattice represent small distortions from its hexagonal parent structure. However in a practical sense it was not obvious whether the risk would be realized or what symptoms would be expected. Therefore we used the following method to predict the unit cell splitting that would be observed if some lattices were misindexed.

Some inconclusive evidence was given by the histogram of *β* angles in the monoclinic cluster, where a bimodal peak is apparent (Figure 5, noted as *P* 2_1_ alternate). Supposing that this peak represented two alternative indexing senses, we plotted the *a* and *c* axes against the *β* angle (Figure 7). One of the clusters is clearly the *P* 6_3_ setting, as it is near 120°. The other peak is split into two sub-peaks, and for these, *a* is similar for both, but *c* is slightly shorter for the smaller value of *β*. For the sake of argument, we propose that the denser of the two sub-clusters corresponds to a single indexing sense; thus we read a set of lattice parameters (*a*, *c*, and *β*) from the center of the denser sub-cluster. We find *a* = 281.02 Å, *c* = 285.83 Å, *β*=119.33°. (We will see that the length of *b* is not important in this argument.)

**Fig. 7.**
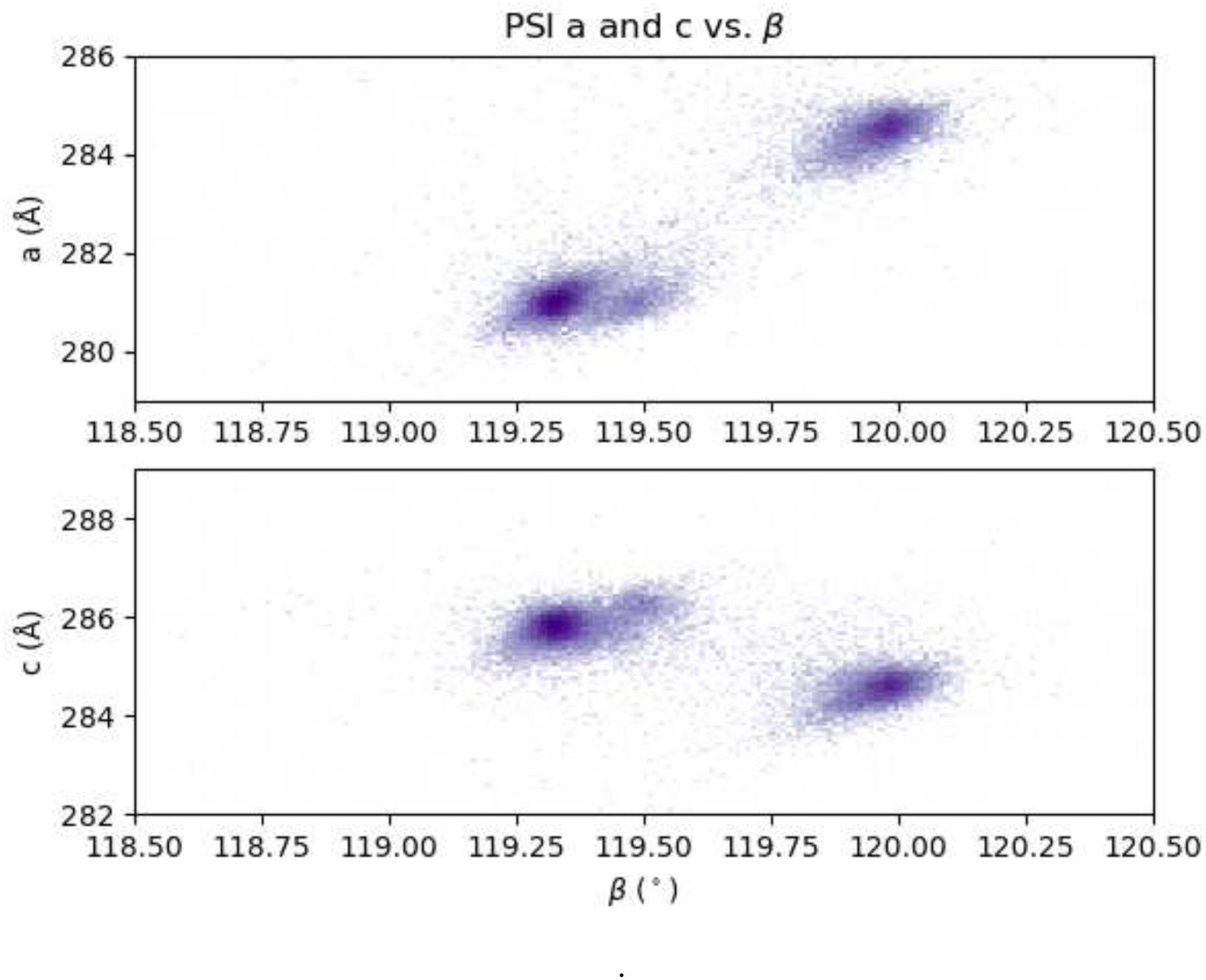
*a* and *c* axes vs. *β* for PSI cells indexed in *P* 2_1_

To illustrate the monoclinic ambiguity more clearly, we draw the monoclinic unit cell in projection along the *b*-axis and label the *a* and *c* axes (Figure 8a). Then we observe that the axis −*a* − *c* is also a lattice vector and therefore could be (mis-) identified as a basis vector in an indexing step. For convenience we label −*a* − *c* as a decoy basis vector *e*. Using simple vector calculations, we find the length of *e* and the two missing angles between *a*, *c*, and *e*. These parameters are labeled on the diagram.

**Fig. 8.**
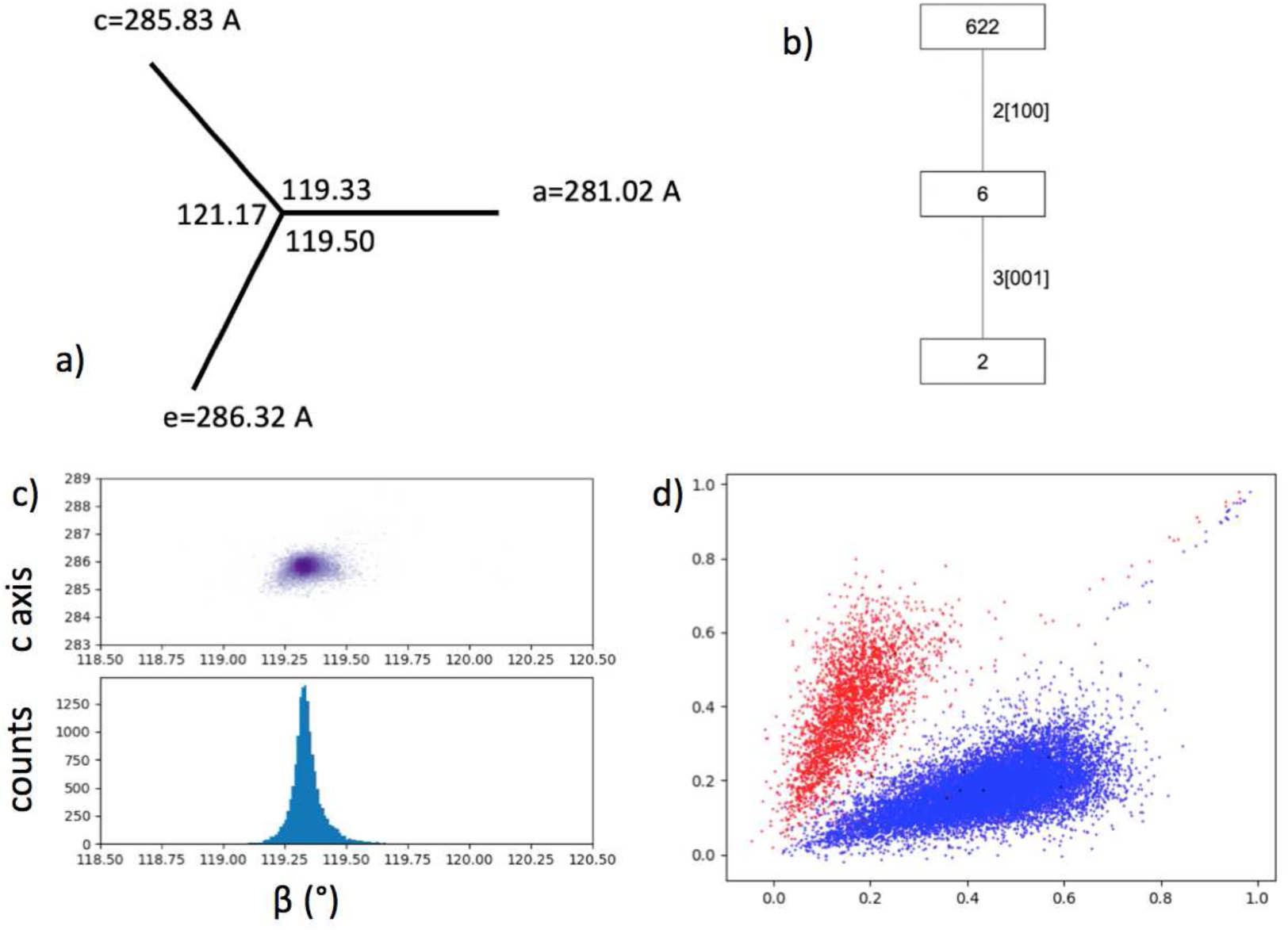
Monoclinic PSI reindexing. a) Diagram of *a* vs. *c* in vector space, with −*a* − *c* labeled as *e*. b) Group-subgroup hierarchy for point group 622. c) *c* vs. *β* after *dials.cosym* using a twofold rotation around the *a* axis. d) The two indexing ambi-guity algorithms (*reindex to reference* and *cosym*) are compared. Correlations of each image to the reference data set using the two indexing operators are shown as *x* vs. *y*, such that the higher of the two would be chosen by *reindex to reference*, and the *cosym* assignments are shown as red vs. blue for the same images.

We note that all of the possible reindexings are apparent in this illustration. The group-subgroup hierarchy, only considering proper rotations, is shown in Figure 8b. The twofold axes of point group 622 are rotations around the axes labeled *a*, *c*, and *e* in Figure. 8a. The rotation around *a* exchanges *c* with *e*, etc. The threefold axis of point group 6 is along the viewing axis and makes *a*, *c*, and *e* equivalent with each other. Thus all six possible reindexings *a*′*, b*′*, c*′ can be generated as follows: (1) Select *a*′ from any one of *a*, *c*, or *e*. (2) Select *c*′ from either of the remaining two. (3) Choose *b*′ from *b* or −*b* to make a right-handed basis.

Importantly, the cell parameters for any of the possible reindexings can be read directly from the diagram. For instance, two of the choices have *a*′ = 281 Å and the other four have *a*′ ≈ 286 Å. We observe in the unfiltered unit cell histograms (Fig. 6) that no measured cells have *c*′ ≈ 281 Å or *a*′ ≈ *c*′ ≈ 286 Å. Therefore, we can eliminate four of the six indexing possibilities. The two remaining ones have *a*′ = 281.02 Å, *c*′ = 285.83 Å, *β* = 119.33° (*a*′*, b*′*, c*′ = *a, b, c*) or *a*′ = 281.02 Å, *c*′ = 286.32 Å, *β* = 119.50° (*a*′*, b*′*, c*′ = *a,* −*b,* −*a* − *c*). These two indexing senses are inter-converted by a twofold rotation around the *a* axis. Both of these unit cells are observed, especially apparent in the *β* vs *c* histogram (Figure 7), and therefore rotation around *a* must be tested as a reindexing operator.

In the existing *dials.cosym* implementation of the Brehm & Diederichs algorithm, the reindexing operators are selected by coset decomposition of the lattice group with respect to the crystal point group (choosing the lattice group with a large tolerance to accommodate pseudosymmetry). In this case, this approach would unnecessarily test all six operators relating point groups 622 and 2. Therefore we implemented a set of parameters for the user to specify one or more reindexing operators as axes and rotations. When a 2-fold rotation around *a* was selected, the resulting *β* histogram and *c* vs. *β* scatter plot were collapsed to a single distribution centered on the expected values of 285.83 Å and 119.33° (Figure 8c). Thus we successfully resolved the pseudomerohedral indexing ambiguity for monoclinic Photosystem I.^8^

Note that both `reindex_to_reference` and `cosym` are able to favorably resolve this ambiguity, as seen in Figure 8d where both algorithms resolve into the same clusters. Generally however, we find that cosym is more reliable for ambiguity resolution.

## 9. Small unit cell chemical crystallography

Finally, a note on small molecule chemical crystallography using *cctbx.xfel*. We recently showed it is possible to solve XFEL structures from materials with tiny unit cells, including just a few angströms on a side (Schriber *et al*., 2022). These images can be indexed using as few as 3 reflections using a graph theory algorithm described in Brewster *et al*. (2015). These *ab initio* structures demonstrate that XFELs are a viable means for material science discovery.

Processing data using this algorithm is done using the program *cctbx.small cell process* and is available using the *cctbx.xfel* GUI. For details please see the *cctbx.xfel* user manual.

## 10. Conclusion

*cctbx.xfel* is a large collaboration and has been used to solve critically important structures at all XFEL facilities. It is under active development and well is supported by government agencies. 100% of *cctbx.xfel* is open source and available on GitHub public repositories. Contributions are welcome.

## 11. Appendix A

Paired refinement was performed on 3 datasets: P450 enzyme Cyp121 resting state (PDB 8TDQ, resolution 1.65Å, Nguyen *et al*. (2023)), Methyl-Coenzyme M Reductase (MCR) oxidized state (PDB 7SUC, resolution 1.90Å, Ohmer *et al*. (2022)), and hen egg-white lysozyme (PDB 7BHK, resolution 1.45Å, Butryn *et al*. (2021)). All datasets were cut at around 10× multiplicity. For Cyp121, higher resolution data (out to 1.5Å) was integrated and merged from deposited raw data in CXI.DB for paired refinement (CXI.DB accession number 222). For each dataset, 5 independent trials of paired refinement were run in each bin and the R-factors are the average R-factors from each of the 5 trials. Both equal width bins and equal volume bins were tested. The paired refinement results for equal width binning (shown here) were more stable than for equal volume binning, perhaps because equal width binning means at higher resolution more reflections are used per bin, allowing more robust sampling of weaker data.

A custom script for replicating this procedure is available at https://github.com/asmit3/eden/blob/main/refinement/paired refinement/paired refinement.py.

## 12. Acknowledgments

We’d like to thank Johannes Blaschke, Robert Bolotovsky, Nicholas Devenish, Gwyn-daf Evans, Vidya Ganapati, Richard Gildea, Johan Hattne, James Holton, Artem Lyubimov, Takanori Nakane, Lee James O’Riordan, Vanessa Oklejas, James Parkhurst, David Waterman, Graeme Winter, Felix Wittwer, and Peter Zwart for their many conversations and contributions to all aspects of *cctbx*, *cctbx.xfel*, and *DIALS*. We’d also like to acknowledge Christopher J. Ohmer and Stephen W. Ragsdale for the their contribution of MCR data and Moritz Kretzschmar, Miao Zhang, Jan Kern, Junko Yano and Vittal K. Yachandra for their contribution of the PSI and PSII data.

This work was supported by an NIH grant R35-GM151988 to NKS, by the US DIALS National Resource, NH grant R24GM154040 to ASB, and the US Department of Energy Integrated Computational and Data Infrastructure for Scientific Discovery grant DE-SC0022215 to ASB.

This research used resources of NERSC, a User Facility supported by the Office of Science, DOE (contract DE-AC02-05CH11231). Use of the LCLS at SLAC National Accelerator Laboratory is supported by the DOE, Office of Science, OBES (contract DE-AC02-76SF00515)

## Footnotes

1 Help for *cctbx.xfel* and DIALS command line programs can be shown using -h, which can be specified multiple times to increase verbosity. Use -v to include parameter descriptions, which can also be spec-ified multiple times to increase verbosity. To see all parameters in detail, run `dials.stills process -c -e10 -a2`

2 Small cell is not currently available in *dials.stills process*, but in the wrapper program *cctbx.xfel.small cell process*. See Schriber *et al*. (2022) for more details.

3 It is not recommended that this cutoff, nor the cutoff for the volume of the envelope estimated from the mosaic parameters around the Ewald sphere, be modified. If crystals are too mosaic or the RMSD is too high, it is likely indicative of an indexing failure or poor starting detector geometry and wavelength.

4 The legacy program, *cxi.merge*, is deprecated.

5 The parameters `min ct=200` and `max ct=300` under `select.significance filter` are recommended for larger unit cells. For small unit cells, leave out these values and use the defaults. This is an unresolved issue in the program.

6 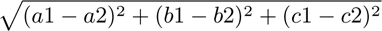

7 The *cctbx.xfel* GUI can make this plot, but in this case *dials.combine experiments* was used to generate a single file with all the indexed crystal models, then *cctbx.xfel.plot uc cloud from experiments* with the parameter `iqr ratio=1.5` was used to generate this plot from the command line.

8 For reference, the specific parameters to enable this were *modify.cosym.twin axis* = 1, 0, 0 and *modify.cosym.twin rotation* = 2

